# In-Cell Synthesis of *N*^ε^-acetyl-L-lysine for Facile Protein Acetylation

**DOI:** 10.1101/2025.10.29.685286

**Authors:** Mengru Zhan, Yanan Zhang, Xiuxiu Ma, Jinyuan Xu, Yinxun Lu, Jia Zhang, Shihan Wu, Jiapeng Zhu, Ren Xiang Tan, Nanxi Wang

## Abstract

N^ε^-acetylation of proteins is a crucial post-translational modification (PTM) that occurs across all kingdoms of life and plays key roles in nearly every cellular process. Due to its broad significance, there is a strong demand for chemical biology tools that enable efficient, site-specific acetylation of target proteins in live cells. Genetic code expansion (GCE) has emerged as a powerful tool for introducing N^ε^-acetylation at specific lysine residues in proteins. However, achieving adequate expression levels typically requires adding 2-10 mM of chemically synthesized N^ε^-Acetyl-L-lysine (AcK) to the culture medium, which can be cumbersome and costly. To overcome this limitation, we present the first proof-of-concept for a one-pot acetylation platform that simplifies protein acetylation in both *E. coli* and mammalian cells. Our approach begins with the discovery of 10 novel lysine acetyltransferases (KATs) capable of biosynthesizing AcK from basic carbon sources. When these enzymes are co-expressed with the genetic incorporation machinery for AcK, they facilitate streamlined, site-specific acetylation of any target protein without compromising *E. coli* viability. This innovative platform not only broadens the range of unnatural amino acids (UAAs) that can be biosynthesized and incorporated but also provides a powerful tool for probing the histone and non-histone acetylation events in live cells. In addition, this technique offers an eco-friendly and scalable method to produce synthetic acetylated proteins, which will provide practical value in acetylation/deacetylation-related research such as chemical biology, biotechnology, and drug development.

## Introduction

N^ε^**-**acetylation of proteins is a highly dynamic and reversible post-translational modification (PTM) that plays a fundamental role in controlling various aspects of cellular function that affect protein functions. These include regulating protein stability, enzymatic activity, subcellular localization and crosstalk with other PTMs, as well as controlling protein–protein and protein–DNA interactions^1^. Understanding specific protein acetylation events is crucial for elucidating their regulatory roles and involvement in disease development, which can, facilitate the development of therapeutics.

To produce N^ε^**-**acetylation of recombinant proteins, the unnatural amino acid N^ε^-Acetyl-L-lysine (AcK) has been chemically synthesized and incorporated into proteins via genetic code expansion^2-3^. This approach has provided valuable tools for studying the acetylation of both histones and non-histone proteins ^4-5^. Efficient AcK incorporation often requires the addition of 2-10 mM of AcK^4, 6-7^. However, the AcK amount that can be uptake by host cells is far less, which hinders its potential in vivo studies. To address this limitation, we first examined whether esterification^8^, a well-established pro-drug technique known for enhancing drug cellular uptake, could facilitate AcK incorporation and thereby reduce its required usage. As illustrated in **Supplementary Figure S1**, the incorporation efficiency of the ethyl-esterified form of AcK (AcK-OEt) increased by approximately twofold compared to its unmodified form, suggesting that higher intracellular AcK levels can significantly enhance its incorporation. This finding motivates us to further explore alternative strategy to enhance intracellular AcK level in order to facilitate AcK incorporation.

Given that intracellular biosynthesis of AcK from basic carbon sources can directly bypass the uptake process, we hypothesized that this approach could effectively promote the expression of synthetic acetylated proteins. Indeed, biosynthesis of UAAs has proven to be an effective solution to the low incorporation efficiency associated with UAAs suffer from cell impermeability. For instance, the poor uptake of negatively charged UAAs, such as phosphothreonine (pThr)^9^ and sulfo-tyrosine (sTyr)^10^ has been circumvented by in-cell biosynthesis catalyzed by heterologously expressed enzymes, allowing for exploration of protein PTMs. Recently, similar strategies have been employed to engineer autonomous bacterial cells capable of biosynthesizing and genetically incorporating polar UAAs, including p-amino-phenylalanine (pAF)^11-12^, p-nitro-L-phenylalanine (pN-Phe)^13^, 5-hydroxyl-tryptophan (5HTP)^14^, 3,4-dihydoxy-L-phenylalanine (DOPA)^15^, and a wide array of UAAs containing aromatic thiol motifs^16^. Despite these exciting advances, most genetically encodable biosynthesized UAAs are analogues of aromatic amino acids, with lysine analogues receiving less attention. To date, D-Cys-*ε*-Lys, facilitating the creation and selection of cyclized therapeutic peptides, is the only lysine-derived UAA that has been successfully biosynthesized and incorporated^17^. The discovery or establishment of novel biosynthetic pathways for additional lysine-derived UAAs, especially AcK, will undoubtfully expand our toolbox of genetic code expansion for protein acetylation studies.

Here, we report a one-pot acetylation platform that seamlessly integrates the biosynthesis and genetic incorporation machineries of AcK into both *E. coli* and mammalian cells. Using bioinformatics analysis, we identified and characterized 10 distinct lysine acetyltransferases (KATs) from various species capable of biosynthesizing AcK in *E. coli*. Protein crystallography and site-directed mutagenesis provided insights into the molecular basis of AcK biosynthesis, while RNA-seq analysis confirmed that AcK biosynthesis has minimal impact on *E. coli* viability. Furthermore, we demonstrated the facile acetylation of non-histone proteins in mammalian cells and histones in *E. coli* cells, paving the way for future investigations into numerous acetylation events and their biological significance in live cells.

## Results

### Identification of lysine acetyltransferases for N^ε^-Acetyl-L-lysine biosynthesis

In nature, KATs catalyze the transfer of the acetyl groups from acetyl-CoA to the ε-amine groups of lysine residues in proteins^18^. To date, three major KAT families, including GCN5/PCAF, p300/CBP, and MYST, have been extensively characterized due to their essential roles in modulating cellular processes^1, 19^. These enzymes have been reported to almost exclusively acetylate specific lysine residues of histones or non-histone proteins, rather than cytoplasmic lysines (**Figure 1a**). Hence, it remains a mystery whether there are KATs that can catalyze the formation of N^ε^-Acetyl-L-lysine (AcK) via acetylation of the ε-amine group of cytoplasmic lysines (**Figure 1b**). To identify KATs capable of introducing N^ε^-acetylation to free lysines inside cells, we first examined *Ce*SSAT, an N-acetyltransferase from *Caenorhabditis elegans*, which is known to be capable of catalyzing the N^ε^-acetylation of lysine in vitro, although its favorable substrates are thialysine and O-(2-aminoethyl)-L-serine^20^.

**Figure 1.**
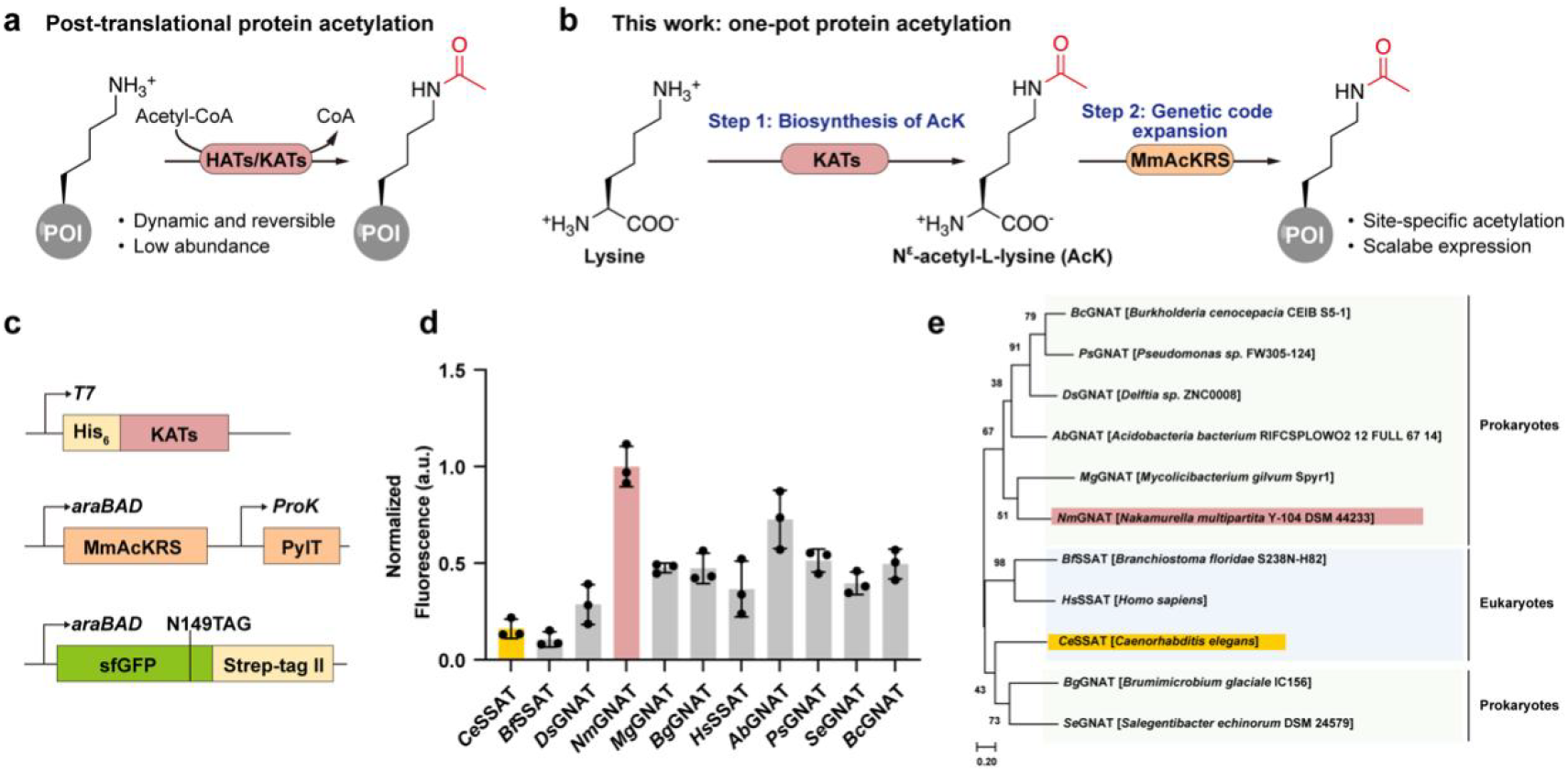
Identification of lysine acetyltransferases for N^ε^-Acetyl-L-lysine biosynthesis. (a) Schematic diagram of a typical post-translational protein acetylation process. (b) Schematic diagram showing one-pot protein acetylation by integrating biosynthesis and genetic encoding of AcK into proteins of interest. (c) Fluorescence-based assay facilitating the tracking of biosynthesized AcK in *E. coli* cells. (d) Screening of lysine acetyltransferases for efficient AcK biosynthesis with fluorescence-based assay. Data are presented as the means of normalized fluorescence from n=3 independent samples. a.u. stands for arbitrary unit. (e) Phylogenetic tree highlighting the the relatedness of lysine acetyltransferases capable of producing AcK among different species. All listed enzymes have been examined in **d**. The enzyme with pink label is *Nm*GNAT, originated from *Nakamurella multipartita* Y-104 DSM 44233. The enzyme with yellow label is *Ce*SSAT, originated from *Caenorhabditis elegans*.

To evaluate the acetyltransferase activity of *Ce*SSAT towards lysine, the gene encoding *Ce*SSAT was codon-optimized, appended with N-terminal His_6_ tag, and cloned into pRSFDuet vector. To simplify tracking of biosynthesized AcK, we developed a fluorescence-based assay in *Escherichia coli* (**Figure 1c**). If AcK can be biosynthesized, pEvol-*Mm*AcKRS encoding the evolved *Methanosarcina mazei* pyrrolysyl-tRNA synthetase (*Mm*AcKRS)/tRNA^Pyl^ pair can specifically recognizes AcK and then suppress the amber stop codon of pBad-sfGFP-149TAG and produce green fluorescence^21-22^. The co-transformants of the above three plasmids in *E. coli* BL21 (DE3) were cultured in LB medium at 30 °C for 16 h without supplying exogenous AcK, and then their relative fluorescence unit (RFU) were recorded. As expected, cells expressing *Ce*SSAT produced detectable green fluorescence, indicating successful biosynthesis and incorporation of AcK in *E. coli* cells (**Figure 1d**).

We envisioned that enzymes with moderate sequence similarity to *Ce*SSAT might retain its catalytic capability while displaying a stronger substrate preference for lysine, thereby enhancing AcK biosynthesis activity. To verify our hypothesis, we performed sequence similarity network (SSN) analysis using *Ce*SSAT as the input sequence to fully explore the potential candidate enzymes for AcK biosynthesis. Consistent with our hypothesis, most of the 500 enzymes finally obtained belong to acetyltransferases and their sequence similarity with *Ce*SSAT is less than 50%.

These enzymes were clustered into nine groups based on the Joint Genome Institute (JGI) classification of enzyme families, from which ten representative sequences were selected and cloned into pRSFDuet vector (**Supplementary Figure S2**). We examined their catalytic activities by the fluorescence-based assay, and to our delight, full length sfGFP expression was observed in all enzymes tested (**Figure 1e)**. In general, enzymes originated from prokaryotes showed stronger AcK biosynthesis activity than those from eukaryotes. Specifically, cells expressing *Nm*GNAT, a putative GCN5-related N-acetyltransferase from *Nakamurella multipartita*, produced the strongest fluorescence, a 6.2-fold increase over *Ce*SSAT (**Figure 1d**). To further explore AcK biosynthesis, we focused on comparing *Ce*SSAT and *Nm*GNAT in the following studies.

### Genetically encoded acetylation via AcK biosynthesis in *Escherichia coli*

After identifying *Nm*GNAT as a promising candidate for AcK biosynthesis, we validated its use of acetyl-CoA and lysine, as demonstrated by the *in vitro* enzymatic assay with purified *Nm*GNAT (**Supplementary Figure 3**). Next, we quantified the intracellular AcK concentrations in *E. coli* cells either by *Nm*GNAT-mediated biosynthesis or exogenous feeding. After 8 h of induction, the cellular AcK concentration accumulated to 6.9 mM, which is 2.5 times higher than that of cells fed with 5 mM AcK, a concentration commonly used in recombinant protein expression (**Supplementary Figure 4**).

To enhance AcK biosynthesis, we screened several culture conditions. We first evaluated the effect of feeding AcK precursors using the forementioned fluorescence-based assay. When 5 mM sodium acetate or 2 mM lysine was supplemented to the LB medium, the green fluorescence of cells expressing *Nm*GNAT reached a peak (**Figure 2a, 2b**). When additional amounts of sodium acetate and lysine were provided, the fluorescence dropped dramatically, indicating that appropriate intracellular levels of precursor molecules have a significant impact on the production and genetic encoding of AcK. Next, we assessed whether culture temperature affects sfGFP149AcK expression and found that cells exhibited the strongest fluorescence at 30 °C (**Figure 2c**). We reasoned that at this temperature, AcK produced by *Nm*GNAT and AcK consumed by *Mm*AcKRS/tRNA^Pyl^ might have reached equilibrium. Finally, we screened culture medium for optimal expression of sfGFP149AcK. Among the media examined, Terrific Broth (TB) medium yielded the highest level of green fluorescence, followed by LB medium and 2xYT medium, likely due to its rich sources of nutrients and AcK precursors (**Figure 2d**). Interestingly, even with a low concentration of glucose at 0.5% w/v in LB medium, the expression of sfGFP149AcK was noticeably impaired. This suggests that the carbon catabolite repression (CCR) effect of glucose may have been more prominent than its role as a carbon source in promoting cell growth and AcK production^23^. Considering the ease of experimentation and protein yield, we conducted expression in LB medium at 30 °C for 16 h in the subsequent experiments, unless specified otherwise.

**Figure 2.**
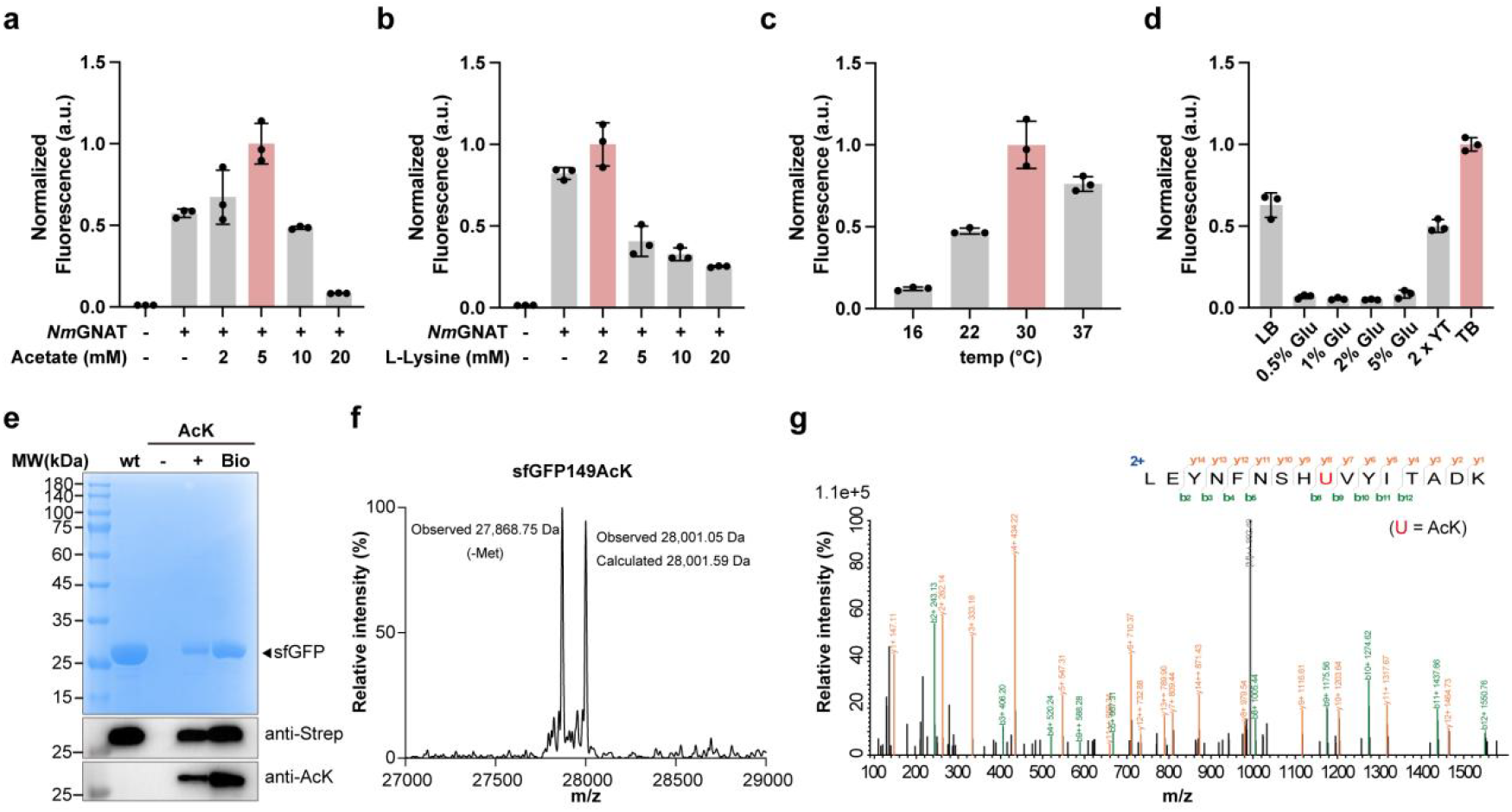
Genetically encode biosynthesized AcK into proteins in *E. coli* cells. (a-d) Screening for optimal conditions for AcK biosynthesis: concentrations of acetate (a) and lysine (b), culture temperature (c), and culture medium (d). (e) SDS-PAGE and western blots showing sfGFP149AcK expression in the absence (-) or presence (+) of 5 mM exogenous AcK or induction of *Nm*GNAT expression (Bio). (f) ESI-TOF MS spectrum of intact sfGFP149AcK expressed with biosynthesized AcK. (g) Tandem MS spectrum of sfGFP149AcK showing precise incorporation of biosynthesized AcK.

We set out to validate the incorporation specificity of biosynthesized AcK into proteins in *E. coli*. The pBad-sfGFP-149TAG reporter was co-expressed with pEvol-*Mm*AcKRS in BL21 (DE3). In the absence of AcK, full-length sfGFP was not detected by Coomassie blue stain or western blot against Strep-II tag. When AcK was exogenously supplied at 5 mM or biosynthesized via *Nm*GNAT expression, full-length sfGFP149AcK was produced at yields of 0.54 mg/L and 2.58 mg/L, respectively (**Figure 2e**). Notably, the latter achieved 37.6% sfGFP expression compared to the wild type sfGFP under the same culture conditions. Western blots against AcK further confirmed the incorporation of AcK into sfGFP and demonstrated that *Nm*GNAT is a robust producer of AcK in *E. coli*.

We then analyzed the purified sfGFP149AcK containing AcK biosynthesized by *Nm*GNAT using electrospray ionization time-of-flight mass spectrometry (ESI-TOF MS) (**Figure 2f**). The spectra showed two peaks: one peak measured at 28,001.05 Da corresponds well to the intact mass of sfGFP149AcK (calculated mass: 28,001.59 Da), and a second peak measured at 27,868.75 Da corresponds to sfGFP149AcK lacking the initiating Met (calculated mass: 27,869.20 Da). Next, to ensure the accurate incorporation of AcK, we performed LC-MS/MS analysis and verified that sfGFP-149 was the only residue carrying acetylation (**Figure 2g**). The data collectively showed successful biosynthesis of AcK, which was incorporated into the specified position of sfGFP-149TAG. Similar to *Nm*GNAT, *Ce*SSAT also enabled efficient biosynthesis and precise incorporation of AcK into sfGFP-149TAG, achieving a lower expression yield of 1.12 mg/L (**Supplementary Figure 5**).

### Structural insights into AcK biosynthesis catalyzed by *Ce*SSAT and *Nm*GNAT

To elucidate the mechanism of AcK biosynthesis, we attempted to crystallize *Nm*GNAT and *Ce*SSAT, both alone and in complex with the substrates, lysine and acetyl-CoA, as well as in complex with the products AcK and CoA. However, only apo*Ce*SSAT successfully crystallized; all other attempts failed. The overall structure of *Ce*SSAT is assembled as a dimer in the asymmetric unit, and the structure of each monomer is similar to that of a typical GCN5-related *N*-acetyltransferase (GNAT) (**Figure 3a**). Inspired by apo*Ce*SSAT, we hypothesized that apo*Nm*GNAT also exists and functions in the form of dimers (**Figure 3b**). The alignment of apo*Ce*SSAT with the AlphaFold3-predicted structure of apo*Nm*GNAT indicates a high structural similarity, with a TM score of 0.9 and an RMSD of 1.69, despite their low sequence identity of 36% (**Figure 3c**). Multiple sequence alignment of *Ce*SSAT with other representative KATs also showed highly conserved secondary structures, suggesting that they are evolutionarily and functionally conserved (**Supplementary Figure 6**).

**Figure 3.**
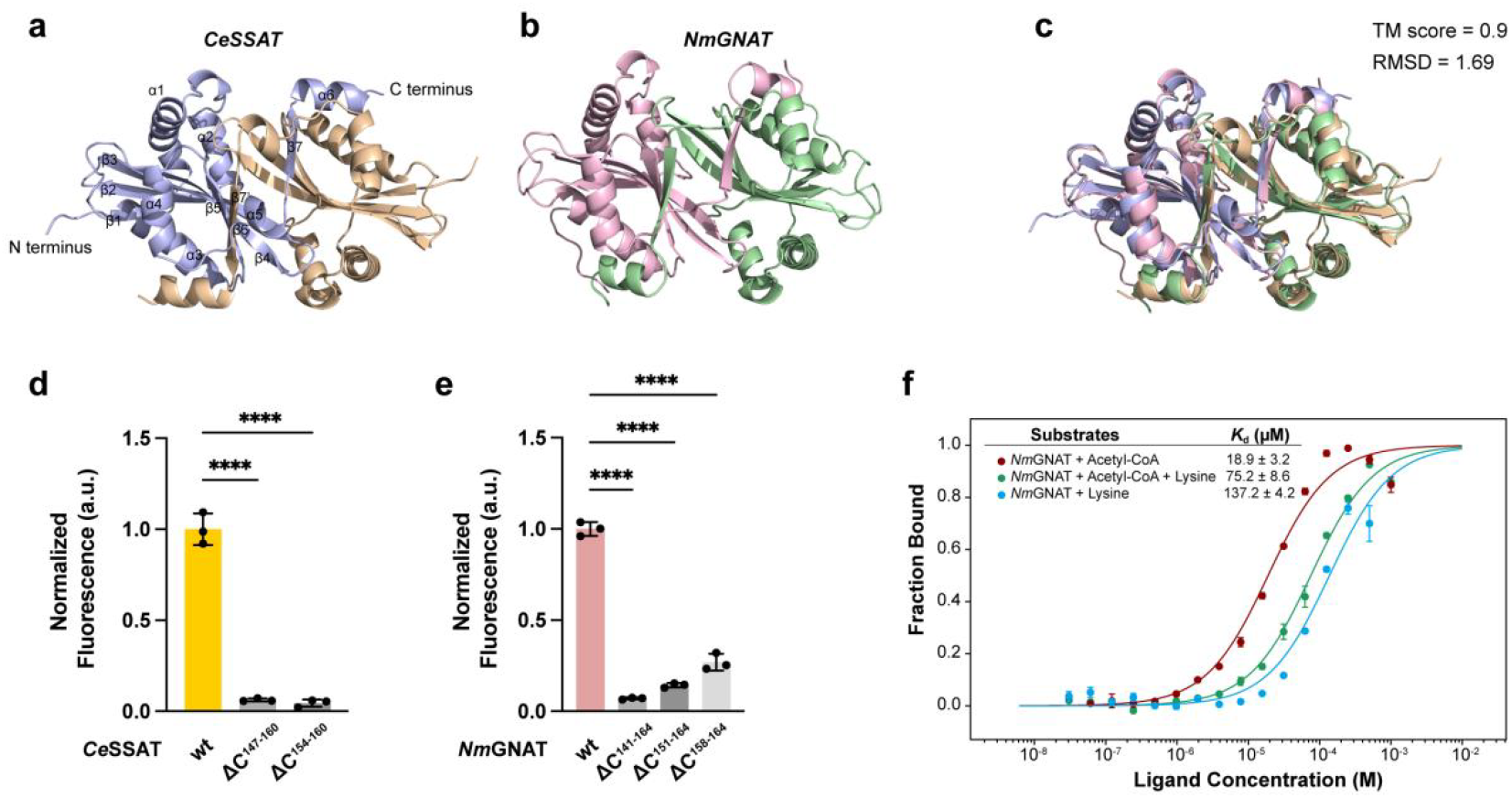
Uncovering the structural basis of *Ce*SSAT and *Nm*GNAT-catalyzed AcK biosynthesis. (a) The crystal structure of *Ce*SSAT (PDB ID: 8YV7). (b) The crystal structure of *Nm*GNAT predicted by AlphaFold3. (c) Superimposition of *Ce*SSAT and *Nm*GNAT highlighting significant structural similarity. (d) Fluorescence-based assay with wildtype *Ce*SSAT (wt) or *Ce*SSAT without the C-terminus residues 147-160 (ΔC^147-160^) or 154-160 (ΔC^154-160^). (e) Fluorescence-based assay with wildtype *Nm*GNAT (wt) or *Nm*GNAT without the C-terminus residues 141-164 (ΔC^141-164^) or 151-160 (ΔC^151-164^) or 158-164 (ΔC^158-164^). (f) MST analysis showing a moderate binding affinity between *Nm*GNAT and its substrates, acetyl-CoA and lysine.

Based on apo*Ce*SSAT structure, its dimer formation involves the C-terminal arm of each monomer: strand β7’ of the opposing monomer consists of residues F145-D150 and extends between β6 and α5, forming a β sheet together with β1 to β6 (**Figure 3a**). To explore whether the C-terminal arms are essential structural motifs, we examined the enzymatic activities of C-terminally truncated *Ce*SSAT and *Nm*GNAT mutants. Cleavage of residues 147-160 or 154-160 resulted in a dramatic decrease in the activity of *Ce*SSAT to produce sfGFP-149AcK by 93.8% and 95.4%, respectively (**Figure 3d**). Similarly, cleavage of residues 151-164 or 158-164 in *Nm*GNAT also led to an obvious reduction in sfGFP production by 85.8% and 73.1%, while a longer truncation of residues 141-164 expanded the reduction to 93% (**Figure 3e**). These results unambiguously demonstrated the contribution of C-terminal arms to AcK biosynthesis through dimer formation.

To explore the residues involved in *Ce*SSAT substrate binding, we conducted protein-ligand docking using Protein-Ligand Interaction Profiler (PLIP) and LigPlot^+^ v2.2.8. The docking structures indicated that E28, L84, I86, R91, N123, N125 and Y130 are involved in acetyl-CoA binding, while S72, T73, E82, D83, R114 and R148 might interact with lysine (**Supplementary Figure 7**). To verify the roles of these residues to *Ce*SSAT activity, we introduced alanine mutations at each site individually and compared their activities by quantifying the resulting fluorescence changes (**Supplementary Figure 8**). We found that the E28A, Y130A, S72A, D83A, and R114A mutations reduced *Ce*SSAT activity to below 50%, whereas the other mutations had minimal or no impact.

To determine the general importance of these mutations, we transferred them to homologous sites into *Nm*GNAT. We found that the transferred mutations, E30A, Y136A, S77A, D89A, and R120A indeed resulted in a significant loss of activity, indicating their broad importance (**Supplementary Figure 8**). Three additional mutations, T78A, E88A, and R151A also decreased *Nm*GNAT activity, likely due to interference with lysine binding. In general, *Nm*GNAT is more sensitive to single mutations than *Ce*SSAT, especially at sites associated with lysine binding. This is likely because lysine is *Nm*GNAT’s preferred substrate, whereas it is not for *Ce*SSAT.

Finally, we determined the binding affinity of *Nm*GNAT and its substrates using a MicroScale Thermophoresis (MST) assay. The *K*_*d*_ of *Nm*GNAT with lysine and acetyl-CoA was 75.2 ± 8.6 μM, indicating moderate binding strength and affinity. The *K*_*d*_ of *Nm*GNAT with lysine alone was 137.2 ± 4.2 μM, while with acetyl-CoA alone it was 18.9 ± 3.2 μM, suggesting a much stronger binding affinity towards acetyl-CoA.

### Transcription responses to AcK biosynthesis in *E. coli*

Next, we performed RNA-Seq to accurately assess the impact of AcK addition or biosynthesis-mediated by *Nm*GNAT on gene expression in *E. coli* cells. Compared to the negative control, the addition of 5 mM AcK led to 809 significant changes (Fold Change≥2, FDR<0.05) in gene expression in BL21 (DE3) cells (**Figure 4a**). Among the 407 downregulated genes, most were primarily involved in metabolic processes, such as the citrate cycle, starch and sucrose metabolism, and pyruvate metabolism (**Supplementary Figure 9**). The remaining 402 upregulated genes were associated with categories related to ribosomes, fatty acid biosynthesis and metabolism, and ABC transporters (**Supplementary Figure 9)**. Notably, when *Nm*GNAT expression was induced, only three genes exhibited significant changes compared to the negative control (**Supplementary Figure 10)**. This suggests that, unlike the addition of AcK, AcK biosynthesis has a minimal effect on gene expression in BL21 (DE3) cells.

**Figure 4.**
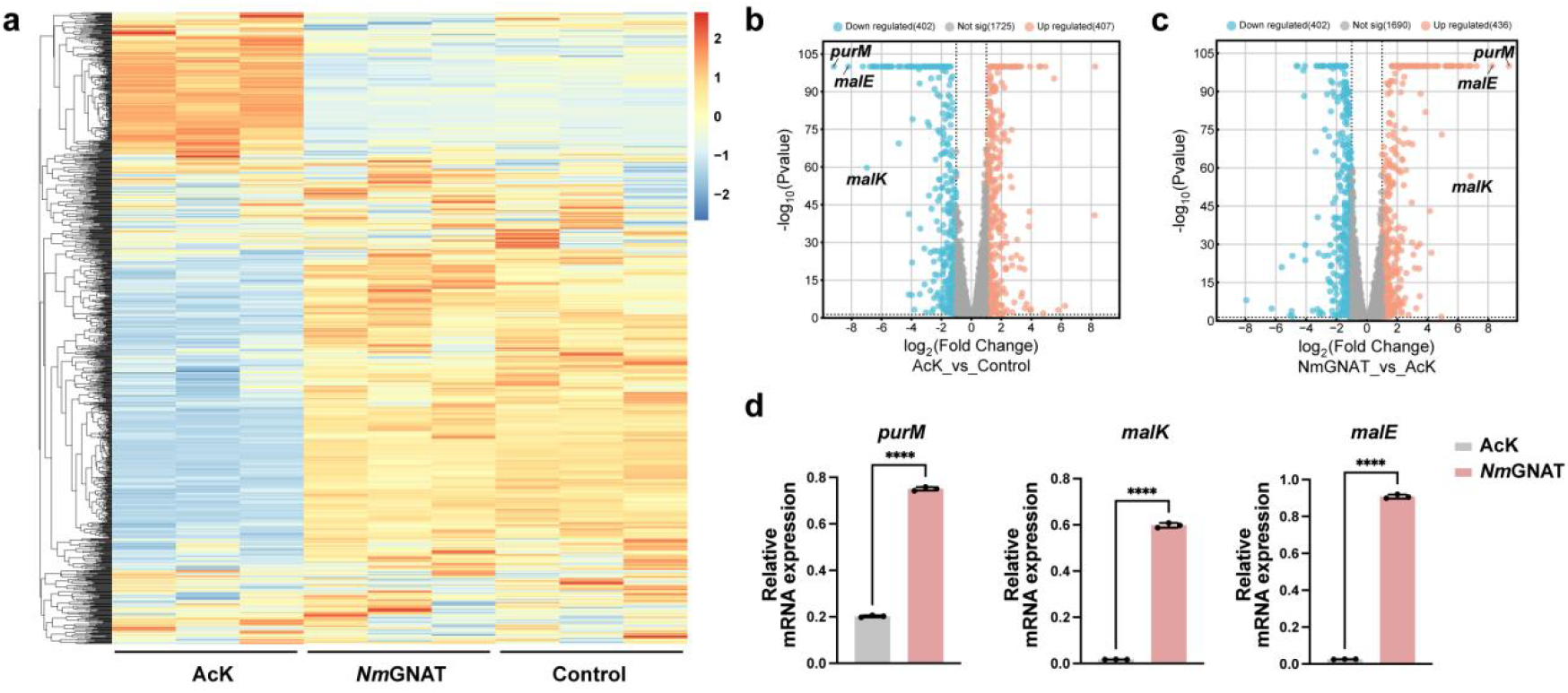
Analysis of AcK biosynthesis-induced transcription responses in *E. coli* cells. (a) RNA-seq analysis of *E. coli* BL21 (DE3) cells treated with 5 mM AcK, expressing *Nm*GNAT, and an empty vector control. (b) Volcano plots illustrating significant transcriptional changes following AcK addition compared to the control. (c) Volcano plots showing distinct transcriptional reponses in BL21 (DE3) cells treated with AcK and expressing *Nm*GNAT. (b-c) Volcano plots were generated using SRplot with a fold change cutoff of 2.0 and a p-value cutoff of 0.05. (d) RT-qPCR analysis of *purM, malK, malE* expression in BL21 (DE3) cells treated with 5 mM AcK or expressing *Nm*GNAT. Data represent the mean ± S.D. (n=3 biological replicates/group) and the p value was calculated by unpaired t-tests. ****p<0.0001.

Among these differentially regulated genes, *purM, malK*, and *malE* caught our attentions because they were significantly downregulated in cells with AcK addition but upregulated in cells expressing *Nm*GNAT (**Figure 4b**). These changes in transcript-level were further confirmed with RT-qPCR analysis (**Figure 4c**). Interestingly, all three genes are involved in essential metabolic processes that support cell growth and energy production. *purM* is crucial for nucleotide biosynthesis, while *malK* and *malE* are vital for carbohydrate metabolism and transport. Changes in their expression reflect the adaptation of cells to different environmental conditions and nutrient availability due to AcK addition or biosynthesis. Collectively, we concluded that AcK biosynthesis has a minor impact on gene expression in *E. coli* cells, fully mimicking the desired metabolic state and energy balance of bacteria, making it a powerful tool for genetically encoding acetylation.

### Genetically encoded acetylation via AcK biosynthesis in mammalian cells

Protein acetylation plays a crucial role in regulating various cellular processes, and is widely recognized for its impact on gene expression, protein stability, and cellular signaling. To achieve efficient protein acetylation in mammalian cells, researchers have developed transient transfection and stable cell line systems based on genetic code expansion. Nevertheless, these methods require the addition of exogenous AcK, which is not convenient for cellular or *in vivo* studies and applications. To streamline protein acetylation, we designed a dual-function plasmid by introducing a self-cleaving peptide P2A between *Ce*SSAT/*Nm*GNAT and *Mm*AcKRS to integrate AcK biosynthesis and genetic encoding machinery (**Figure 5a**). As expected, HEK293T cells expressing *Ce*SSAT and *Nm*GNAT actively produced and accumulated cellular AcK to concentrations of 0.11 mM and 0.19 mM, respectively, 48 hours post-transfection. These levels are slightly lower than those observed in cells supplemented with 1 mM AcK (**Figure 5b**).

**Figure 5.**
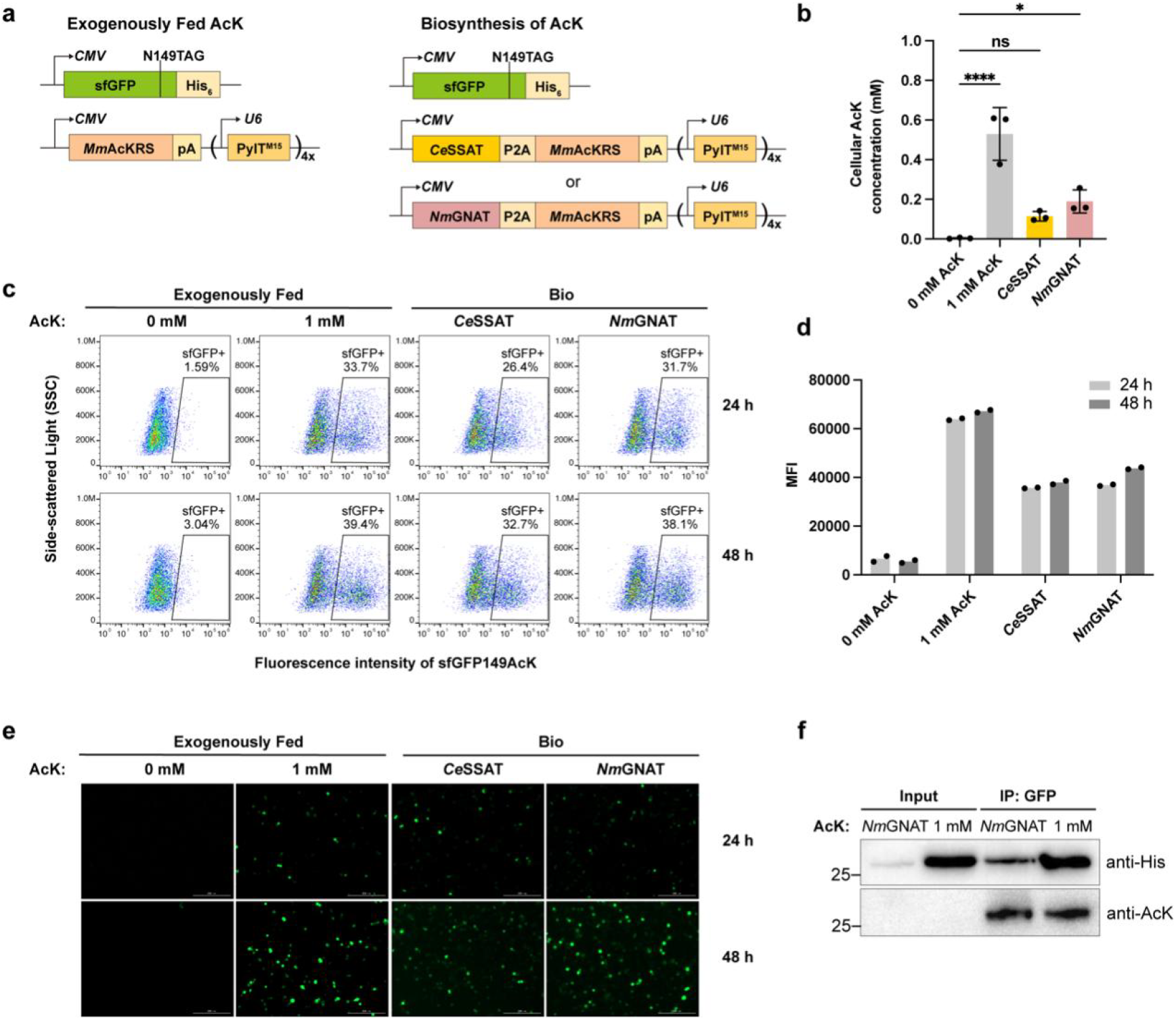
Genetically encoding biosynthesized AcK into proteins in mammalian cells. (a) Schematic diagram of mammalian expression system for generating AcK-containing proteins in the presence of exogenously fed AcK or biosynthesized AcK. (b) Cellular AcK concentrations in mammalian cells treated with 0 mM and 1 mM AcK or when expressing *Ce*SSAT and *Nm*GNAT. Data are plotted as the mean ± standard deviation from n = 3 independent samples and the p value was calculated by one-way ANOVA. *p=0.0359, ****p<0.0001, ns stands for non-significance. (c) Flow cytometry analysis of AcK incorporation into sfGFP149TAG in HEK293T cells treated with exogenous or biosynthesized AcK for 24 h and 48 h. (d) Comparison of AcK incorporation efficiency of the samples in (c). (e) Representative images of HEK293T cells expressing sfGFP149AcK in the presence of exogenous and biosynthesized AcK for 24 h and 48 h. (f) Immunoprecipitation (IP) analysis of sfGFP149AcK produced in HEK293T cells expressing *Nm*GNAT or treated with 1 mM AcK.

To evaluate the incorporation efficiency and fidelity of biosynthesized AcK, the dual-function plasmid encoding *Ce*SSAT/*Nm*GNAT and *Mm*AcKRS/PylT^M15^ pair were co-transfected with the reporter plasmid pcDNA-sfGFP-149TAG^24^. For comparison, *Mm*AcKRS/PylT^M15^ pair was co-expressed with the reporter in the presence or absence of 1 mM exogenously fed AcK. The expression of full-length sfGFP was monitored by flow cytometry and fluorescent microscopy 24- and 48-hours post-transfection. To our delight, transient expression of *Ce*SSAT and *Nm*GNAT achieved efficient production of sfGFP149AcK, comparable to that observed with the addition of 1 mM AcK (**Figure 5c and 5e**). Consistent with the cellular AcK concentrations, the expression levels of sfGFP in cells biosynthesizing AcK were slightly lower than in cells fed with AcK, as indicated by the mean fluorescence intensity (MFI) (**Figure 5d**). We further performed immunoprecipitation (IP) to enrich acetylated sfGFP from HEK293T cells treated with *Nm*GNAT or exogenously fed AcK and confirmed that either approach can efficiently introduce AcK into the designated site of protein of interests (**Figure 5f**). Overall, in mammalian cells, *Nm*GNAT is more efficient than *Ce*SSAT in producing AcK, which is similar to what we observed in *E. coli*.

### Expression of synthetic acetylated histone using the one-pot acetylation approach

Chromatin is a complex mixture of DNA and proteins that responds to external stimuli, such as cellular metabolites and drug treatments, to regulate DNA utilization within cells^25^. Histones, a major component of chromatin, undergo various post-translational modifications (PTMs). Since the first report of histone acetylation in 1964, there is an ever-growing list of PTMs on histones have been identified, including crotonylation, malonylation, succinylation, glutarylation, β-hydroxybutyrylation, dihydroxyisobutyrylation, benzoylation, and most recently, lactylation^26-27^. Although these PTMs can be genetically encoded into target proteins via the GCE^28-32^, this approach requires the addition of chemically synthesized UAAs, which, according to our RNA-seq analysis, may act as external stimuli to disrupt proteome in live cells. We hypothesized that our one-pot acetylation approach could bypass this limitation by enabling seamless incorporation of the biosynthesized AcK, thereby minimizing the potential impacts on cellular processes.

Acetylation of histone H4 at lysine 5 (H4K5ac) is a critical modification that influences chromatin dynamics and gene expression. However, the detailed molecular mechanisms of this modification and its crosstalk with other PTMs remain poorly understood^33-34^. To address this, we used our one-pot acetylation approach to express this acetylated histone in *E. coli* as a proof of concept. We cloned both the wild-type histone H4 gene (H4-wt) and a mutant containing an amber stop codon at the K5 site (H4-K5TAG) into the pTXB1 vector. The C-terminally fused Mxe-CBD domain on this vector allows for the release of the expressed H4 peptide through DTT cleavage. As shown in **Figure 6a**, H4K5ac-Mxe-CDB fusion proteins were expressed and detected using anti-Strep-II tag antibody, both with exogenous addition of AcK and with AcK biosynthesis via induction of *Nm*GNAT. However, the yield of H4K5ac was significantly lower with exogenous AcK addition, achieving only 19.8 % of the yield observed in the biosynthesis group. This low yield explains why the signal from the exogenous AcK group was barely detectable using pan anti-acetyllysine or anti-acetyllysine-H4K5 antibodies, while the biosynthesis group exhibited strong signals (**Figure 6b and 6c**). Importantly, after DTT treatment, the released H4K5ac peptide was detected by dot blot assay (**Supplementary Figure 11**), further validating the utility of this approach in generating acetylated peptides. To this end, we demonstrated the general applicability of our one-pot acetylation approach for investigating both non-histone and histone proteins.

**Figure 6.**
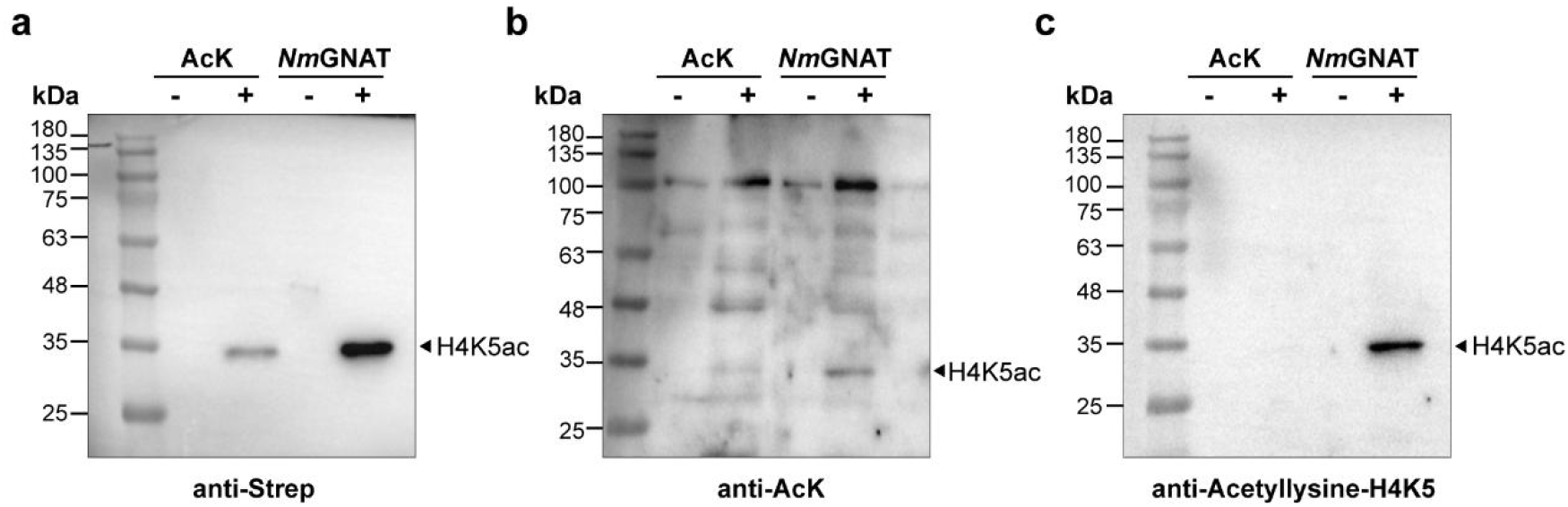
Production of histone H4K5ac using the *E. coli*-based one-pot acetylation method. (a) The expression of full-length H4K5ac was confirmed using ani-Strep antibody to detect the C-terminal Strep tag. Acetylation at the K5 residue was verified with an anti-acetyllysine mouse mAb (b) and a specific anti-acetyl-H4K5ac antibody (c), respectively.

## Conclusions

In summary, we developed a groundbreaking one-pot acetylation approach that integrates biosynthesis with genetically encoded machinery for AcK. This innovative approach streamlines the production of N^ε^-acetylated unnatural amino acid, as well as histone and nonhistone proteins, in an efficient, site-specific, scalable and eco-friendly manner in both *E. coli* and mammalian cells. Given the widespread occurrence and critical role of N^ε^-acetylation in various cellular processes, this approach allows for more detailed and precise investigations into the relationship between specific acetylation events and their biological outcomes, which will offer new insights into cellular regulation and signaling. This approach holds promise for broad and practical applications, such as the production of acetylated antigens for immunological studies and the development of histone deacetylase (HDAC) inhibitors for therapeutic use, as demonstrated by the production of H4K5ac. By expanding the toolbox available for protein acetylation, this work contributes to the fields of chemical biology, biotechnology, and drug development from a new perspective.

## Acknowledgements

This work was supported by the Natural Science Foundation of China (82104050 and 22377058), Specially-appointed professor of Jiangsu Province, and Nanjing University of Chinese Medicine (XPT82104050). Diffraction data of *Ce*SSAT were collected at beamline BL02U1 of the Shanghai Synchrotron Radiation Facility (SSRF, China). The authors thank Dr. Irene Coin in University of Leipzig for generous gift of plasmid pNEU-hMbPylRS-4xU6M15 (Addgene plasmid # 105830). Mr. Tao Pei and Dr. Yan Wu from Nanjing Jiangbei New Area Biopharmaceutical Public Service Platform Co., Ltd. are greatly appreciated for mass spectrometric analysis.

## Author contributions

N.W., R.T., and J.Z. conceived the project. N.W. designed the experiments and analyzed the data. Y.Z. conducted the preliminary study on the discovery of KATs. M.Z., J.X., and Y.L. performed the molecular biological experiments. X.M. performed the protein crystallography analysis. J.Z. carried out the fluorescence imaging experiments. S.W. contributed to chemical synthesis. N.W. and Y.Z. wrote the manuscript.

