## Supplemental File for "In-Cell Synthesis of *N*^ε^-acetyl-L-lysine for Facile Protein Acetylation"

### In-Cell Synthesis of $N^{\varepsilon}$-acetyllysine for Facile Protein Acetylation

#### General experimental procedures

##### Reagent and materials

Molecular biology reagents, including antibiotics such as ampicillin, kanamycin, and chloramphenicol were purchased from Sangon Biotech unless otherwise specified. Restriction enzymes were obtained from New England Biolabs. Molecular cloning and qRT-PCR reagents were obtained from Vazyme. The HRP-conjugated anti-Hisx6 tag antibody (catalog no. HRP-66005) and GFP tag polyclonal antibody (catalog no. 50430-2-AP) were purchased from Proteintech. The anti-acetyllysine Mouse mAb (catalog no. PTM-102) and Anti-Acetyl-Histone H4 (Lys5) Mouse mAb (catalog no. PTM-163) were acquired from PTM Biolabs. The Strep II Tag Mouse Monoclonal Antibody (catalog no. AF2927) was purchased from Beyotime, and the HRP-conjugated Goat Anti-mouse IgG (catalog no. A21010) was obtained from Abbkine.

##### Chemical synthesis of AcK esters

##### Synthesis of methyl $N^{\varepsilon}$-acetyl-*L*-lysinate (AcK-OMe)

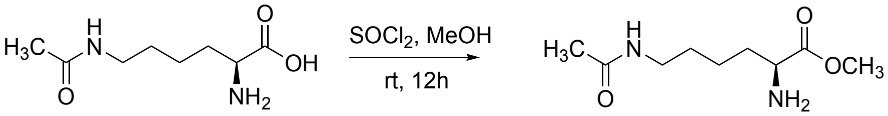

$N^{\varepsilon}$-acetyl-L-lysinate (1.88 g, 10 mmol) was dissolved in 20 mL of MeOH and cooled down to 0 °C. $SOCl_2$ (7.26 mL, 100 mmol) was then added slowly. The reaction was stirred at 25 °C overnight. The solution was subsequently concentrated, and the resulting residue was purified by flash-column chromatography to yield AcK-OMe.

##### Synthesis of ethyl $N^{\varepsilon}$-acetyl-*L*-lysinate (AcK-OEt)

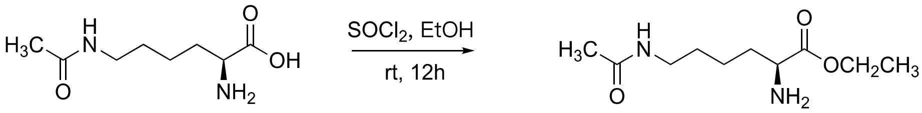

$N^{\varepsilon}$-acetyl-L-lysinate (1.88 g, 10 mmol) was dissolved in 20 mL of EtOH and cooled down to 0 °C. $SOCl_2$ (7.26 mL, 100 mmol) was added slowly. The reaction was stirred at 25 °C overnight. The solution was subsequently concentrated, and the resulting residue was purified by flash-column chromatography to yield AcK-OEt.

##### SSN and phylogenetic analysis of lysine acetyltransferases identified for AcK biosynthesis

We initially created a Sequence Similarity Network (SSN) with JGI-IMG, with *Ce*SSAT from

*Caenorhabditis elegans* as the input sequence. By applying an E-value threshold of 1e-20, we identified a final set of 500 lysine acetyltransferase sequences. These sequences were clustered into nine groups based on SSN analysis, and 11 distinct lysine acetyltransferases from various species were selected for further investigation. We then analyzed the phylogenetic relationships of these enzymes by using their amino acid sequences to generate a phylogenetic tree using the Maximal likelihood method with arithmetic mean by MEGA 11 software package with the bootstrap set at 1000.

**Construction and expression of lysine acetyltransferases**

The genes encoding the 11 identified lysine acetyltransferases were codon-optimized, chemically synthesized and inserted into *Bam*HI-*Hin*dIII sites of pRSFDuet vectors by Tsingkes Biotechnology Co.,Ltd. The amino acid sequences of these enzymes are provided in Table S1. The resulting plasmids were transformed into BL21 (DE3) competent cells. The colonies were grown overnight at 37 °C in LB medium containing 50 μg/mL kanamycin with vigorous agitation. The following day, the cell culture was diluted 1:100 into fresh LB medium and induced at 37 °C for 6 h, with or without 0.25 mM IPTG. The expression of above enzymes was confirmed with SDS-PAGE (data not shown).

**Expression and purification of sfGFP149AcK in *E. coli* cells**

Plasmids pBad-sfGFP-149TAG-StrepII, pRSFDuet-*Nm*GNAT, and pEvol-*Mm*AcKRS were co-transformed into BL21 (DE3) cells and grown on plates containing 100 μg/mL ampicillin, 50 μg/mL kanamycin, and 50 μg/mL chloramphenicol. The colonies were grown overnight at 37˚C in LB medium supplemented with the appropriate antibiotics. The following day, the culture was diluted 1:100 into fresh LB medium and induced with 0.2% w/v arabinose, 0.25 mM IPTG, and 25 mM nicotinamide (NAM) until OD600 reached 0.4~0.6. After induction at 30 ˚C overnight, the proteins were purified using STarm Streptactin prepacked column (Smart-Lifesciences) following the manufacturer's instructions.

**Microscale thermophoresis (MST) experiments**

MicroScale Thermophoresis (MST) was performed to evaluate the biomolecular interactions between *Nm*GNAT and its substrates, lysine and acetyl-CoA. A 100 µL solution containing

100 nM of 6× His-tagged *Nm*GNAT protein in 1× phosphate-buffered saline with Tween detergent (PBST) was mixed with 100 μL of 100 nM Monolith His-Tag Labeling Kit RED-tris-NTA 2nd Generation dye (NanoTemper Technologies, catalog no. MO-L018). The mixture was incubated in the dark at room temperature for 30 minutes to label *Nm*GNAT with fluorescein isothiocyanate (FITC). Lysine and acetyl-CoA, each at a concentration of 4 mM, were then added to the 25 nM protein-dye mixture. The samples were mixed thoroughly, loaded into Monolith standard capillaries (NanoTemper Technologies, catalog no. MO-K025), and their binding affinities were measured using the NT.115 Monolith instrument with Nano-RED excitation and medium MST power settings. Dissociation constants were calculated based on a binding model, with data processed using NanoTemper's Affinity analysis software to ensure precise determination.

**Protein crystallization and structural elucidation**

Purified apo *Ce*SSAT proteins were crystallized using the sitting-drop vapor-diffusion method in MRC-2Drops 96-well plates. Briefly, the apo *Ce*SSAT proteins were mixed with the gel filtration buffer (20 mM Tris-HCl, 150 mM NaCl, pH 8.0) to prepare the crystallization droplets. Then the crystallization drops were incubated at 22 °C against the well solutions (40 mM potassium dihydrogen phosphate, 16% w/v polyethylene glycol 8,000 and 20 % v/v glycerol). The finest plate-like crystals emerged and reached their final size after about one week.

**RNA-seq analysis of BL21 (DE3) cells for AcK biosynthesis**

BL21 (DE3) cells were transformed with either pRSFDuet-*Nm*GNAT (*Nm*GNAT group) or the pRSFDuet empty vector (AcK group or negative control group) and grown on LB-agar plates containing 50 μg/mL kanamycin overnight at 37 ˚C. The resulting colonies were inoculated into seed cultures, which were then diluted 1:100 into LB media containing 50 μg/mL kanamycin and incubated at 37 ˚C. When the OD600 reached 0.1~0.2, the cells were induced with 0.2 mM IPTG, and the AcK group was supplied with 5 mM AcK. Subsequently, the cells were moved to 30 ˚C and harvested when the OD600 reached 0.5~0.6. Transcriptome sequencing was conducted by Tsingkes Biotechnology Co.,Ltd.

To confirm the mRNA expression level, total RNA was extracted from above samples using

FreeZol Reagent (Vazyme, catalog no. R711) and Bacteria RNA Enhancement Reagent (Vazyme, catalog no. R412-C5) according to the manufacturer's instructions. RNA concentration was measured using Nanodrop, and reverse transcription was performed using HiScript II Q RT SuperMix for qPCR (Vazyme, catalog no. R222-01). qRT-PCR was performed with ChamQ Blue Universal SYBR qPCR Master Mix (Vazyme, catalog no. Q312) and the following primers:

malK-F：TCCTGCCGGTAAAAGTGACC

malK-R：GGCGAATACCCAGCGACATA

malE-F：GGGTCTGACCTTCCTGGTTG

malE-R：GATGGTCATCGCTGTTTCGC

gapA-F：GCTCGTAAACACATCACCGC

gapA-R：AGCGTTGGAAACGATGTCCT

purM-F：ATCAGGCTGTTCACTGGTGG

purM-R：GTCGCTGACTTTAGAGCCGT

**Fluorescence imaging and flow cytometric analysis of HEK293T cells expressing sfGFP149AcK**

Approximately $0.1\times10^6$ HEK293T cells were seeded into 12-well plates and cultured for 24 h. The cells were co-transfected with 0.75 μg of pcDNA-sfGFP-149TAG and 0.75 μg pNEU-*Nm*GNAT-P2A-*Mm*AcKRS (*Nm*GNAT group), pNEU-*Ce*SSAT-P2A-*Mm*AcKRS (*Ce*SSAT group) or pNEU-*Mm*AcKRS (AcK group) respectively, using PolyJet transfection reagents (SignaGen Laboratories, catalog no. SL100688) according to the manufacturer's instructions. After 18 h, the culture media were replaced with fresh complete medium containing 1× Deacetylase Inhibitor Cocktail (Beyotime, catalog no. P1112) with or without 1 mM AcK. Then the cells were cultured for an additional 24 h and 48 h. To visually observe the sfGFP149AcK expression, the cells were imaged using BioTek Cytation 5 Cell Imaging Multimode Reader (Agilent). To quantify the incorporation efficiency and fidelity of biosynthesized AcK, the cells were transfected following the same protocol and then collected for flow cytometric analysis using Gallios Flow Cytometer (Beckman Coulter). The mean fluorescence intensity (MFI) was analyzed with Flowjo.

**Expression and immunoprecipitation of sfGFP149AcK in mammalian cells**

To genetically encode biosynthesized or exogenous AcK into target proteins, approximately $3.5\times10^6$ HEK293T cells were seeded into a 10-cm dish and cultured for 24 h. The cells were co-transfected with 5 μg of pcDNA-sfGFP-149TAG and either 5 μg pNEU-*Nm*GNAT-P2A-*Mm*AcKRS (*Nm*GNAT group) or pNEU-*Mm*AcKRS (AcK group) using PolyJet transfection reagents (SignaGen Laboratories, catalog no. SL100688), following the manufacturer's instructions. After 18 h, the culture media were replaced with fresh complete medium containing 1× Deacetylase Inhibitor Cocktail (Beyotime, catalog no. P1112), with or without 1 mM AcK, and the cells were cultured for another 48 h. The transfected cells were then collected by centrifugation at 4 °C, lysed with ice-cold RIPA buffer containing protease inhibitors, 1× Deacetylase Inhibitor Cocktail and centrifuged to remove cell debris. The supernatant was incubated with GFP tag polyclonal antibody (Proteintech, catalog no. 50430-2-AP) at 4 °C overnight with gentle agitation. Subsequently, rProtein A/G MagPoly Beads (Smart-Lifesciences, catalog no. SM037001) were added to the antibody-lysate complex and washed according to the manufacturer's instructions. The eluted proteins were then subjected to western blot analysis with anti-acetyllysine mAb (PTM Biolabs, catalog no. PTM-102) and HRP-conjugated anti-Hisx6 tag antibody (Proteintech, catalog no. HRP-66005).

**Construction and expression of pTXB1-H4-wt and pTXB1-H4K5TAG**

The wild-type histone H4 (H4K5-wt) and the mutant H4 with an amber stop codon at K5 (H4K5TAG) genes were PCR amplified using the following primers and cloned into pTXB1 vector (New England Biolabs, catalog no. N6707S) via homologous recombination, resulting in the plasmids pTXB1-H4-wt-strep and pTXB1-H4K5TAG-strep, respectively. Subsequently, pTXB1-H4K5TAG-strep was transformed into BL21 (DE3)-pEvol-*Mm*AcKRS chemical competent cells under three conditions: 1. with pRSFDuet-*Nm*GNAT (*Nm*GNAT group); 2. without pRSFDuet-*Nm*GNAT but with exogenous AcK (AcK group); 3. without pRSFDuet-*Nm*GNAT and AcK (negative control group). As an expression control, pTXB1-H4-wt-strep was transformed into BL21 (DE3)-pEvol-*Mm*AcKRS chemical competent cells. The tested groups were examined with or without the addition of inducers (0.2% Arabinose and 0.25 mM IPTG) and 30 mM NAM after overnight induction at 30 °C.

H4-insert-F: tttaagaaggagatatacatATGTCAGGACGCGGCAAA

H4-insert-R: agtgcatctcccgtgatgcaCACCTTGCGGTGGCGCTT

pTXB1-H4-vec-F: TGCATCACGGGAGATGCAC

pTXB1-H4-vec-R: ATGTATATCTCCTTCTTAAAGTTAAACAAAAT

**Dot blot and western bot analyses of recombinant histones H4**

*E. coli* cells expressing H4K5-wt and H4K5ac were collected by centrifugation at 4 °C, 12,000 rpm for 10 minutes. The cell pellets washed with 1 mL of PBS and centrifuged agin under the same conditions. The cell pellets were then resuspended in 100 μL of BugBuster protein extraction reagent (Millipore, catalog no. 70584-M) and incubated at room temperature for 10 minutes with gentle agitation. The total proteins were extracted and the soluble fractions were collected by centrifuge at 4 °C, 12,000 rpm for 10 minutes. For dot blot analysis, 2 μL of each protein sample was dotted onto a 0.2 μm nitrocellulose membrane and then dried for 30-60 minutes at 37 °C. For western blot analysis, about 5-10 μL of each protein sample was loaded onto an SDS-PAGE gel, followed by membrane transfer. The membranes were blocked with 5% skim milk, then incubated with anti-acetyllysine Mouse mAb (PTM biolabs, catalog no. PTM-102) and Anti-Acetyl-Histone H4 (Lys5) Mouse mAb (PTM biolabs, catalog no. PTM-163) to detect AcK incorporation, and with the Strep II Tag Mouse Monoclonal Antibody (Beyotime, catalog no. AF2927) to detect the expression of full-length histone H4.

**Mass spectrometric analysis**

The intact protein mass was determined using ESI-MS on a Q Exactive-HF-X mass spectrometer (Thermo Fisher) operating in a positive ESI mode and connected to a Vanquish liquid chromatography unit (Thermo Fisher). The mass spectra of acetylated protein were analyzed with BioPharma Finder (Thermo Fisher). For tandem mass spectrometry, the trypsin-digested peptides were separated by an Easy LC 1200 liquid chromatography unit

(Thermo Fisher) and analyzed with a QE-HF-X mass spectrometer (Thermo Fisher). The mass spectra of acetylated peptide were analyzed with pFind 3.

**Supplementary Tables**

**Table S1. Amino acid sequences of lysine acetyltransferases in this work**

| Enzyme names | Organisms | Amino acid sequences |
|---|---|---|
| *Ce*SSAT | *Caenorhabditis elegans* | MKNFEIVTVTPDHAEQLISMIHELAEFEKMKSSVVN TAEKLRKDIENKAVHGFIAFIGEEPAGMNLFYYAYS TWVGQYLHMEDLYIRPQFRRMGLARTLWKKLAEL ARDKGIVRLEWAVLDWNKNAIALYDTVDYVNLTK SEGWFTFRMDGAAINKFADE |
| *Bf*SSAT | *Branchiostoma floridae* S238N-H82 | MSDFRIRDLQPEDCRELVNMIKELAEFEGLPEQVKI TEETLRRDGFGDRPFYHCLIAEVRNKDSQDGPWLT VGYAMYFYSYGTWVGRMIYLEDLYIKPQYRGKGI GTSMMTKVAQIGVENECQRMQWVVLNWNQGAID FYKKHDSIDLTSDEKWHLFRMEREELRKFATLHDQ TVVKGC |
| *Ds*GNAT | *Delftia* sp. ZNC0008 | MTTTLQIRPATPDDAELIVRFVRELAVYEKAEHEVL ATPEHVKRTLFADNPAVFGLICLHGDQPVGFAVYFF NYSTWQGRHGLYLEDLYVTPEARGLGAGTALLRR LAQIAVEKDCGRFEWSVLDWNLPSIEFYDRLGALP QTEWIRYRVTGDTLLEMAREPA |
| *Nm*GNAT | *Nakamurella multipartita* Y-104, DSM 44233 | MSVPDPRIRPIAPADVPAVVALVHDLAAYEKAPEQ CHLTAAQLHSALFGPSPALFGLIASEPDEPVAGFAL YFLNFSTWEGVHGIYLEDLFVRPQQRGSGLGKALL TRLAQIAVDRGYARVEWSVLDWNTPSIEFYRSLDA VPMNGWTTFRLTGPALRTAAGTG |
| *Mg*GNAT | *Mycolicibacterium gilvum* Spyr1 | MSRRGERSDGGNICIRAVRPGDEAELTAMIHELA EFEHAAEECTVTESRLAQALFGPEPAVYGHIVEV DGQAAATALWFRNYSTWDGVAGIYLEDLFVRPQ FRRRGLGRKLLATLARECVDNGYSRLSWAVLDW NANAIALYDGVGGKPQTEWITYRVSGPGLSELAS RDVPGSGS |
| *Bg*GNAT | *Brumimicrobium glaciale* IC156 | MMEGTNIRKAKRGDELALMGLVHELAEFEKAPDE VINTPEQLAIDIFDDKICDCFVYEIEGIIRGMALYYIS YSTWRGRCLYLEDLYIQPDFRRGGIGQKLFQTLVDE AKEMGVKRMDWQVLDWNDSAIQFYKKIGATLDPE WINGRLFF |
| *Hs*SSAT | *Homo sapiens* | MASVRIREAKEGDCGDILRLIRELAEFEKLSDQVK ISEEALRADGFGDNPFYHCLVAEILPAPGKLLGPC VVGYGIYYFIYSTWKGRTIYLEDIYVMPEYRGQG IGSKIIKKVAEVALDKGCSQFRLAVLDWNQRAM DLYKALGAQDLTEAEGWHFFCFQGEATRKLAGK |

| | | |
|---|---|---|
| *Ab*GNAT | *Acidobacteria bacterium* RIFCSPLOWO2_12_FULL_67_14 | MIADLRIVPATAADAPLLLRLIRDLAEYERLSQDVVATEASLRESLFGAEPGAQAVIAYVGEEPAGVAVWFYNFSTFLGRPGLYLEDLFVRPEWRGRGLGRALLQYLAREAVARRCGRMEWAVLDWNAPAIGFYRSLGAVPMDEWTVYRLTGEALRRLADEERS |
| *Ps*GNAT | *Pseudomonas* sp. FW305-124 | MTIEIRPAVPSDAAQILTFITELAEYEKARHEVIASVVDIERSLFSEGATAHGLICLRDGLPIGFAVFFFSYSTWLGSNCLYLEDLYINPEQRGGGAGKKLLRHLAKIAFDNGCGRFEWSVLDWNEPAIAFYKSIGAQPQEEWVRYRMEGDALRDFAL |
| *Se*GNAT | *Salegentibacter echinorum* DSM 24579 | MNINIRKSTKEDMPAVLELIQELAEFEKEPEAVIISAEDLVRDGFGENPSFTCFVAEVEGKIEGMALCYFRYSTWKGKTVHLEDLVVREKMRGKGLGNALYTRVIEFAKDQGVERAEWVVLDWNKHARDFYQRSGAKVFTNWCTVQMDGPAIEDFLNKGR |
| *Bc*GNAT | *Burkholderia cenocepacia* CEIB S5-1 | MVQIDIRSATVADVPQILRFITELAVYEKAEHEVVATPESLERSLFGEGSPARALMCEIDGEPAGFAVYFFSYSTWLARQGLYLEDLYVSPRFRGAGAGLRLLKALARIAVDSGCGRFEWSVLDWNEPAIRFYESVGAAPQSEWVRYRLAGDELRAFADGTPVSAA |

**Table S2. Diffraction data and refinement statistics for the structure of *Ce*SSAT.**

| **Data statistics** | ***Ce*SSAT** |
|---|---|
| Space group | P 1 |
| Cell dimensions | |
| a, b, c (Å) | 57.33,65.10,65.32 |
| α, β, γ (°) | 80.62,72.76,63.98 |
| Resolution (Å) | 62.32-3.38(3.56-3.38) |
| $R_{merge}$ (%) | 8.2(14) |
| I/σI | 5.4(2.7) |
| Completeness (%) | 89.4 |
| Redundancy | 2.6 |
| **Refinement** | |
| Resolution (Å) | 34.25-3.38(3.50-3.38) |
| No. reflections | 9322 |
| $R_{work}/R_{free}$ (%) | 18.28/25.22 |
| No. atoms | 5261 |
| Protein | 73.30 |
| Ligand/ion | - |
| Water | 71.5 |
| B-factors | 71.45 |
| R. ms. deviations | |
| Bond lengths (Å) | 0.010 |
| Space group | P 1 |
| Cell dimensions | |
| a, b, c (Å) | 57.33,65.10,65.32 |

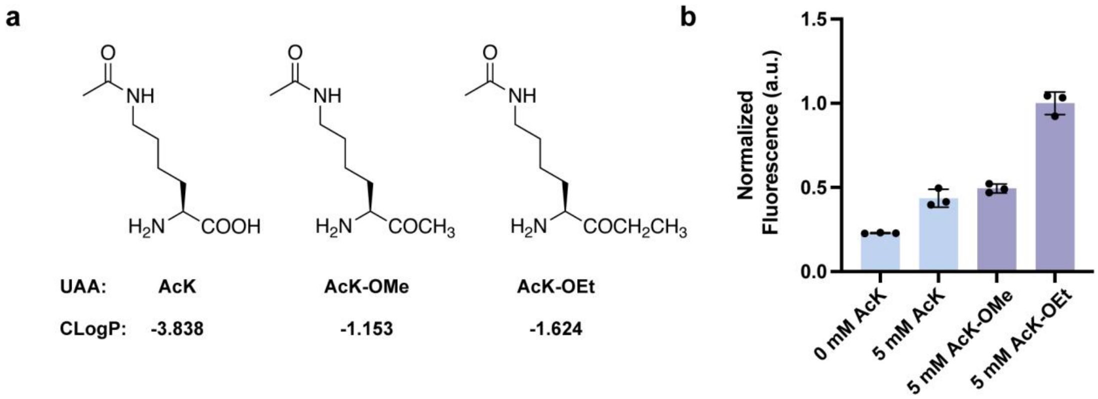

**Figure S1. Expression of sfGFP149AcK in *E. coli* cells using AcK and AcK esters.**

(a) Structures of AcK and AcK esters. (b) The relative fluorescence intensity of cells expressing sfGFP149AcK in the prescence of AcK and AcK esters. **b** Data are plotted as the mean ± SD (n=3 biologically independent samples).

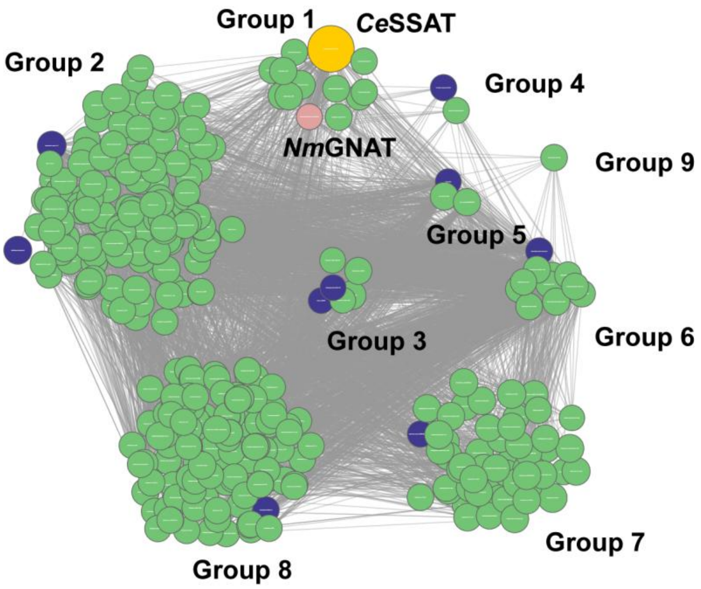

**Figure S2. Identification and clustering of lysine acetyltransferases for AcK biosynthesis using a sequence similarity network.**

Sequence similar network (SSN) was generated by JGI-IMG server using *Ce*SSAT (yellow node) as an input sequence and E value of 20. Each node represents a representative GCN5-related N-acetyltransferase. The size of the node reflects the similarity with the *Ce*SSAT amino acid sequence. The larger the node, the higher the sequence identity of the gene with CeSSAT. Representative nodes in pink and dark blue were selected for gene synthesis and further analysis. The edges indicate the relatedness between the acetyltransferase. (Group1: GCN5-related N-acetyltransferase, Group 2: GNAT superfamily N-acetyltransferase, Group 3: Acetyltransferases, Group 4: diamine N-acetyltransferase, Group 5: spermidine/spermine N1-acetyl transferase 2, Group 6: hypothetical protein, Group 7: L-amino acid N-acyltransferase YncA, Group 8: Ribosomal protein S18 acetylase RimI, Group 9: predicted N-acetyltransferase YhbS.)

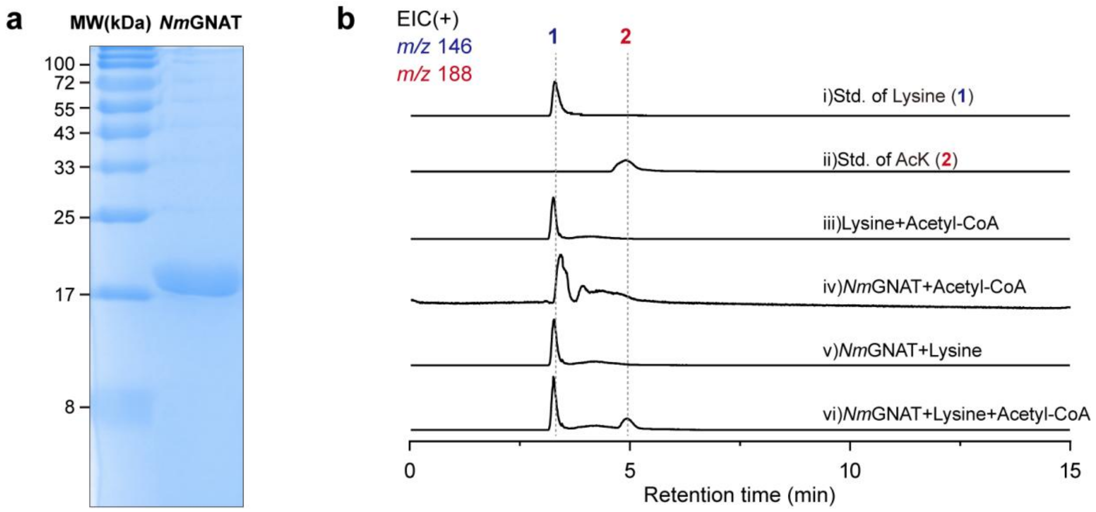

**Figure S3. Biosynthesis of AcK with purified *Nm*GNAT *in vitro*.**

(a) SDS-PAGE of purified *Nm*GNAT. (b) LC-HRMS analysis of AcK production demonstrating that AcK is biosynthesized only in the presence of *Nm*GNAT and its substrates, lysine and acetyl-CoA.

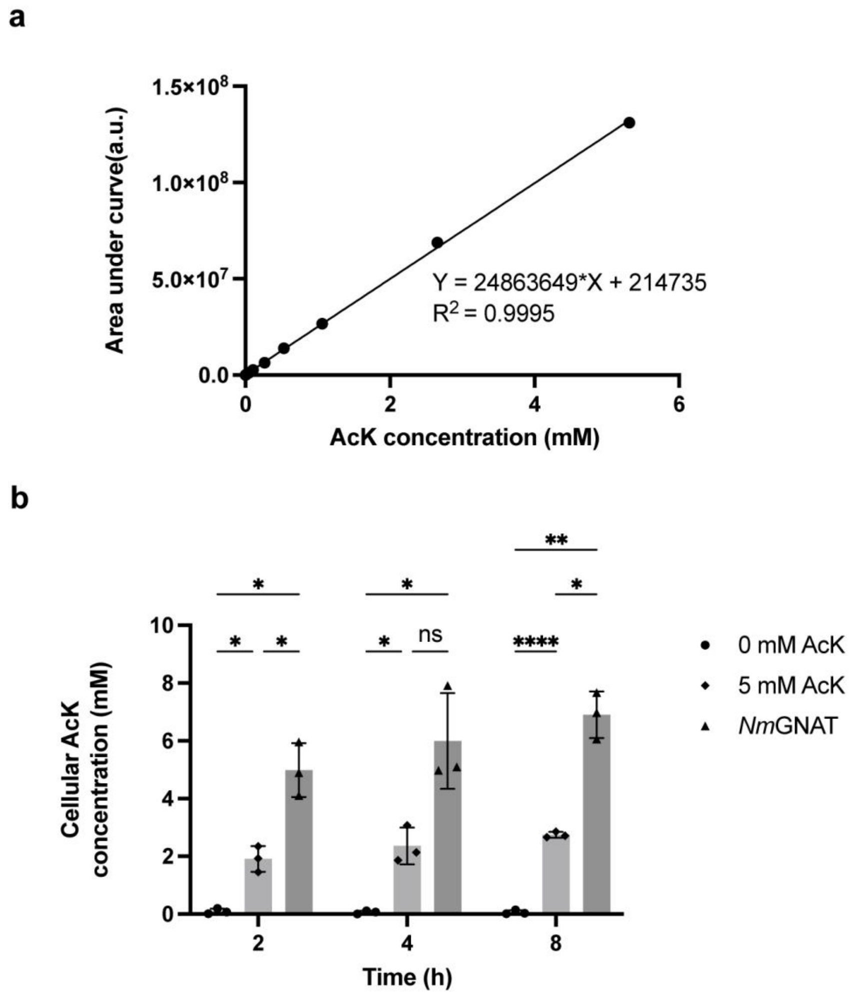

**Figure S4. Measurement of cellular AcK concentrations in *E. coli* cells.**

Standard curve for quantification of cellular AcK concentrations. (b) Concentration of AcK extracted from *E. coli* cell lysates measured at 2, 4, and 8 h after treatment with exogenous AcK or *Nm*GNAT induction. Data are plotted as the mean ± SD (n=3 biologically independent samples). Statistical data were calculated by two-way ANOVA. $^{*}p$=0.0336, $^{**}p$=0.0040, $^{****}p$<0.0001, ns stands for non-significance.

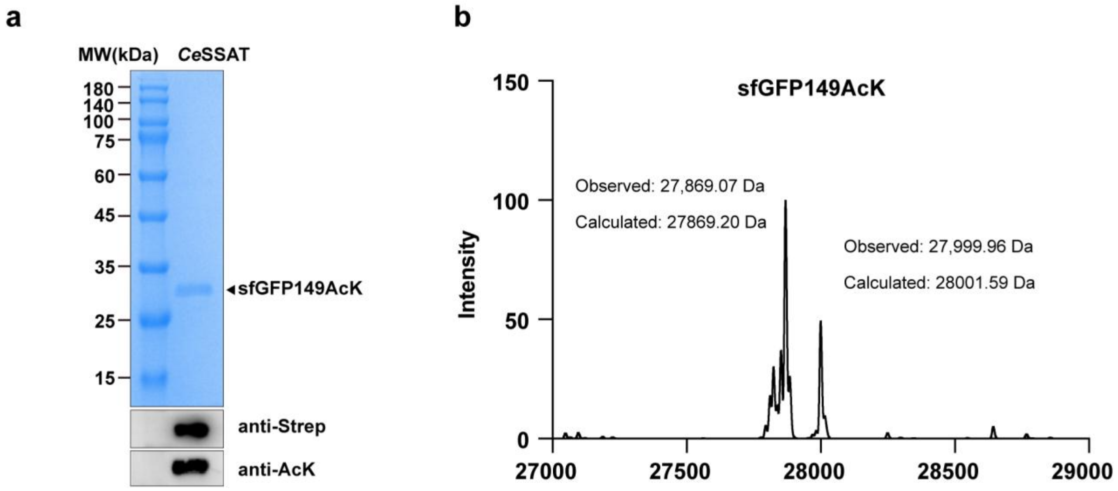

**Figure S5. Genetically encode biosynthesized AcK into proteins in *E. coli* cells with *Ce*SSAT expression.**

(a) SDS-PAGE and western blots showing sfGFP149AcK expression when *Ce*SSAT was induced.

(b) ESI-TOF MS spectrum of intact sfGFP149AcK expressed with biosynthesized AcK using *Ce*SSAT.

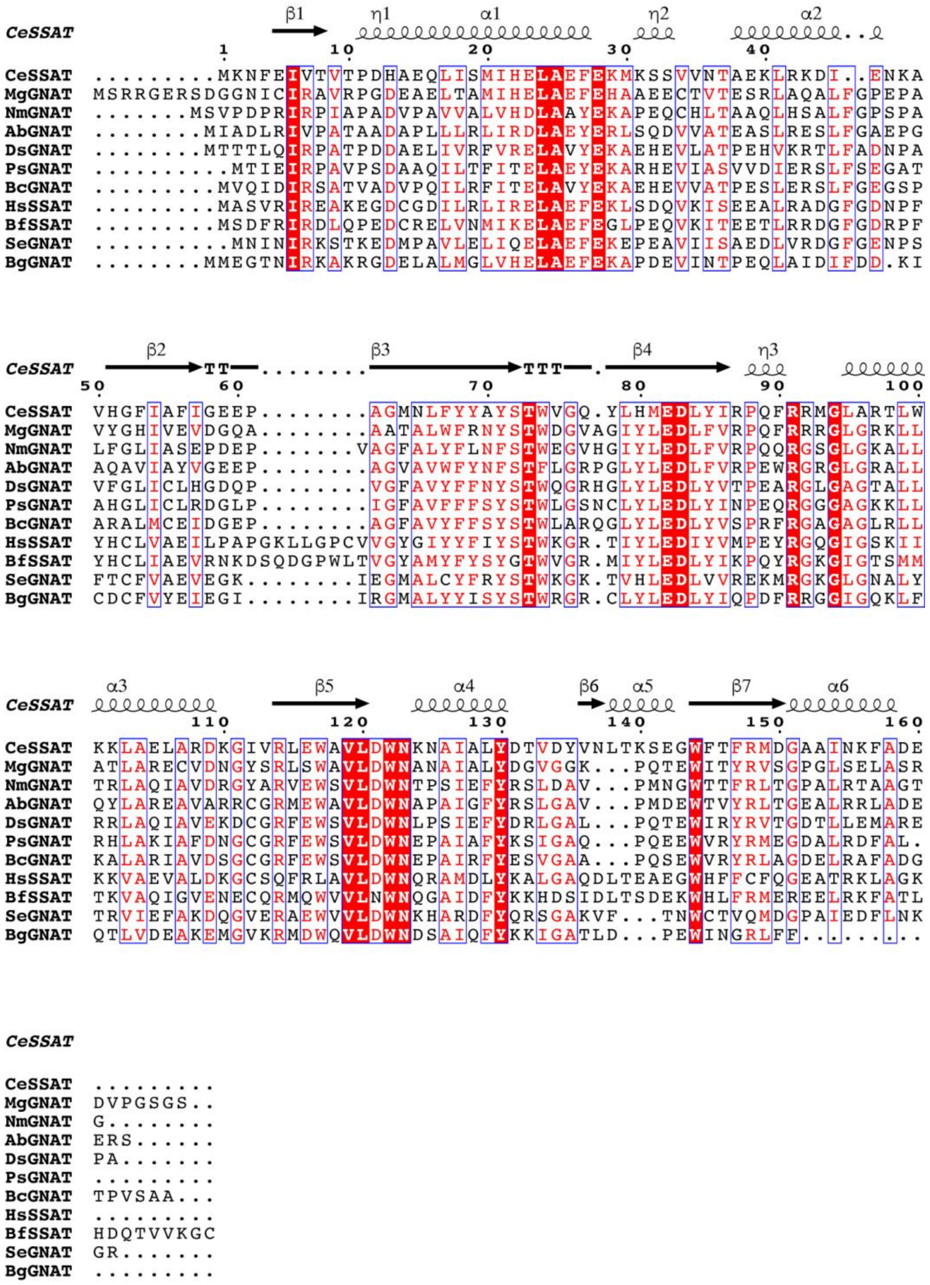

**Figure S6. Multiple-sequence alignment of *Ce*SSAT with 10 other functionally characterized lysine acetyltransferases.**

Multiple sequence alignment (MSA) analysis was performed using CLUSTAL W with the crystal structure of apo *Ce*SSAT as input and visualized using ESPript 3 online program.

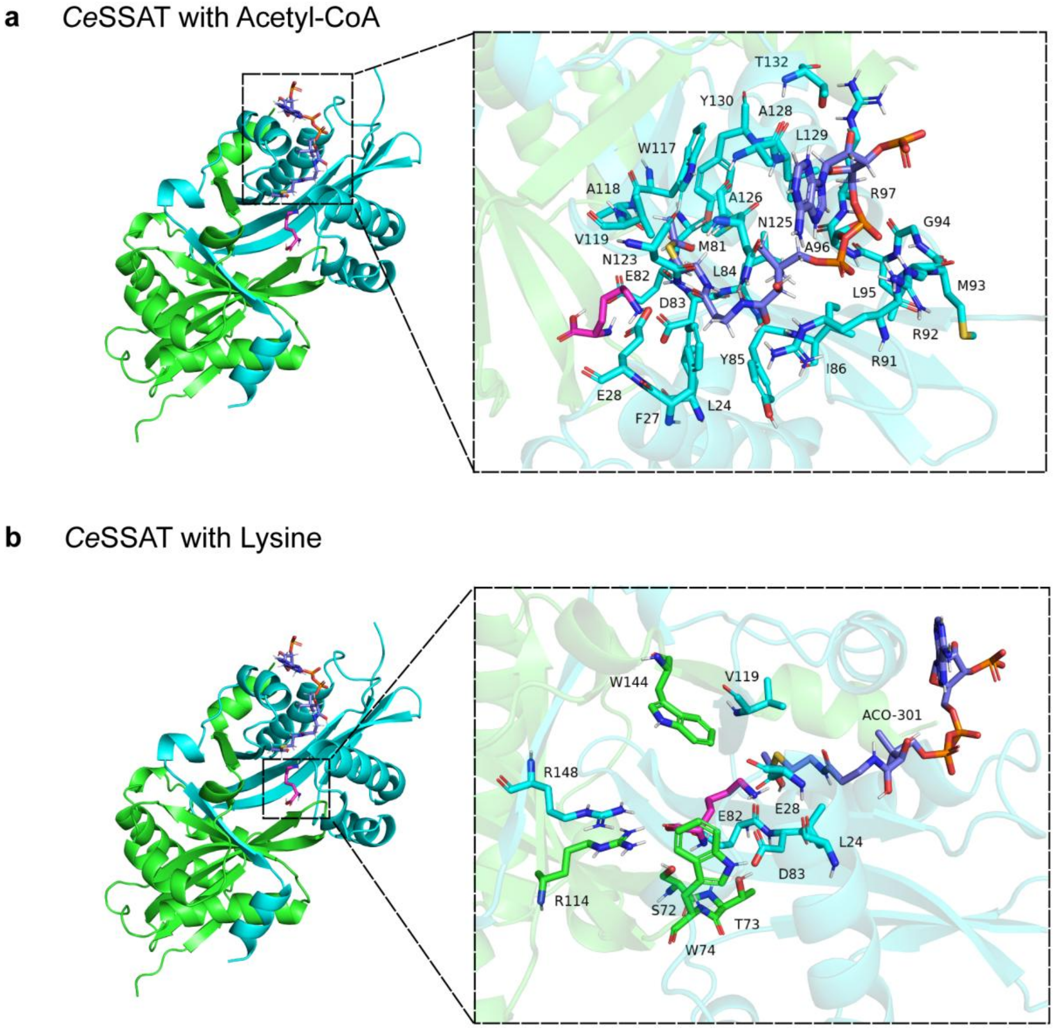

**Figure S7. Docking of *Ce*SSAT structure with acetyl-CoA and lysine.**

The docking structures indicated that (a) E28, L84, I86, R91, N123, N125 and Y130 might be involved in acetyl-CoA (slate) binding, and (b) S72, T73, E82, D83, R114 and R148 might be involved in lysine (magenta) binding.

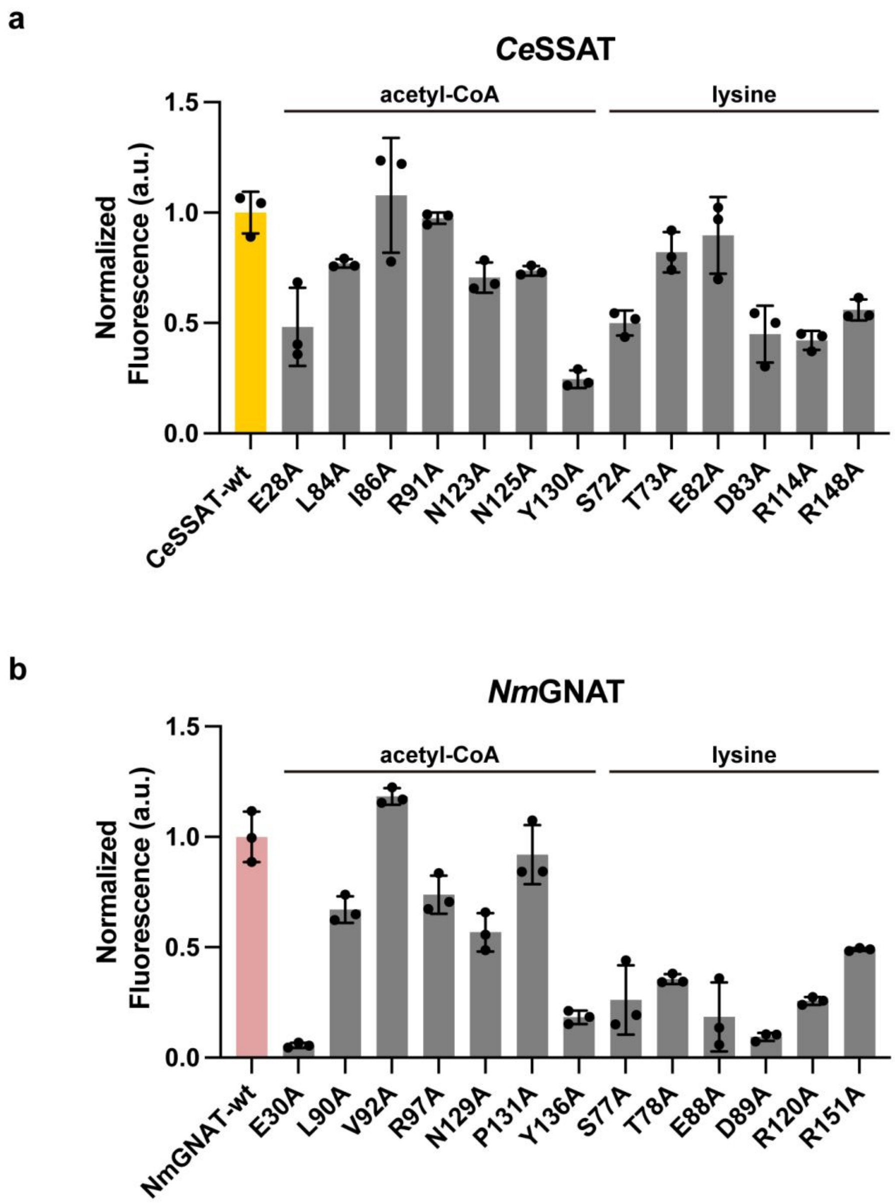

**Figure S8. Site-directed mutagenesis to identify key residues essential for the activity of *Ce*SSAT and *Nm*GNAT.**

(a)Enzyme activities of *Ce*SSAT mutants measured using a fluorescence-based assay. (b) Enzyme activities of *Nm*GNAT mutants measured using a fluorescence-based assay. **a-b** Data are plotted as the mean ± SD (n=3 biologically independent samples).

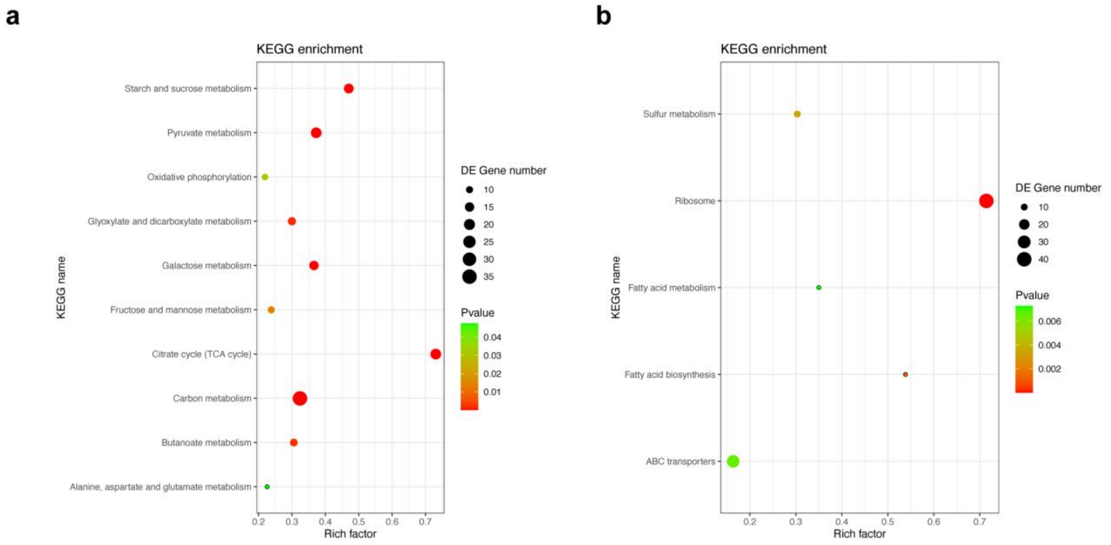

**Figure S9. KEGG enrichment analysis of differentially regulated genes in BL21 (DE3) cells treated with or without AcK.**

(a) Down-regulated genes are mainly associated with the citrate cycle, starch and sucrose metabolism, and pyruvate metabolism. (b) Up-regulated genes are primarily enriched in pathways related to ribosomes, fatty acid biosynthesis and metabolism, and ABC transporters.

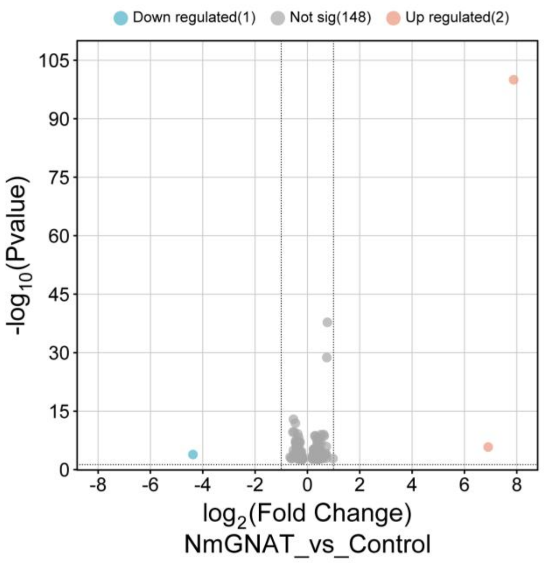

**Figure S10. Volcano plots illustrating transcriptional responses in BL21 (DE3) cells treated with or without *Nm*GNAT induction.**

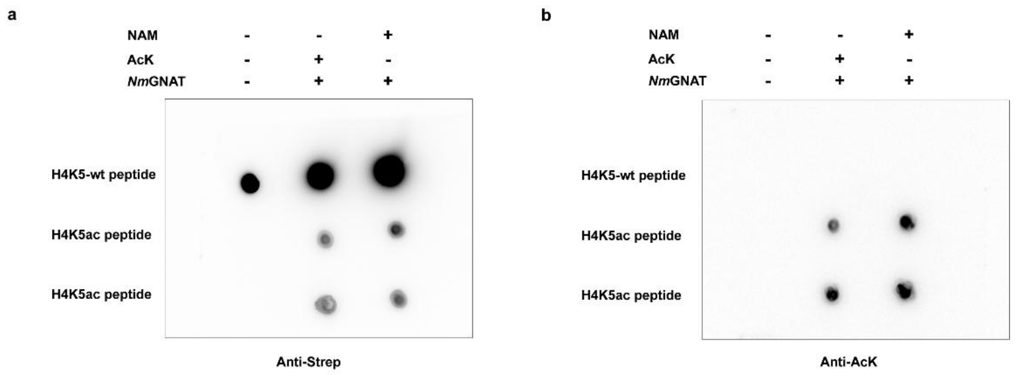

**Figure S11. The H4K5-wt and H4K5ac peptides were detected by dot blot assays.** (a) The expression of full-length H4K5ac was confirmed using ani-Strep antibody to detect the C-terminal Strep tag. (b) Acetylation at the K5 residue was verified with an anti-acetyllysine mouse mAb.

#### Original Images

##### Figure 2 (e)

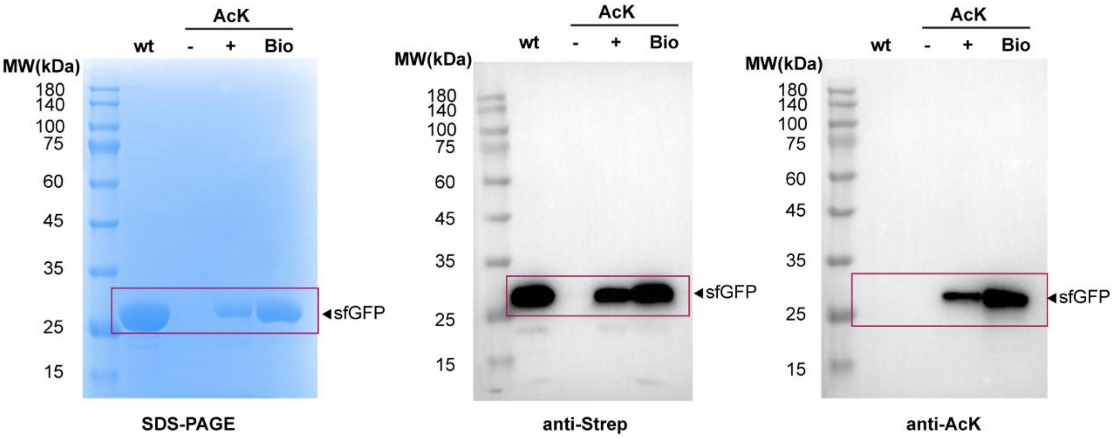

##### Supplement Figure S5 (a)

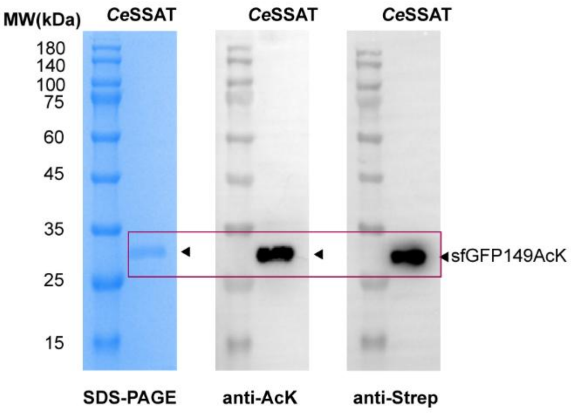

##### Figure 5 (f)

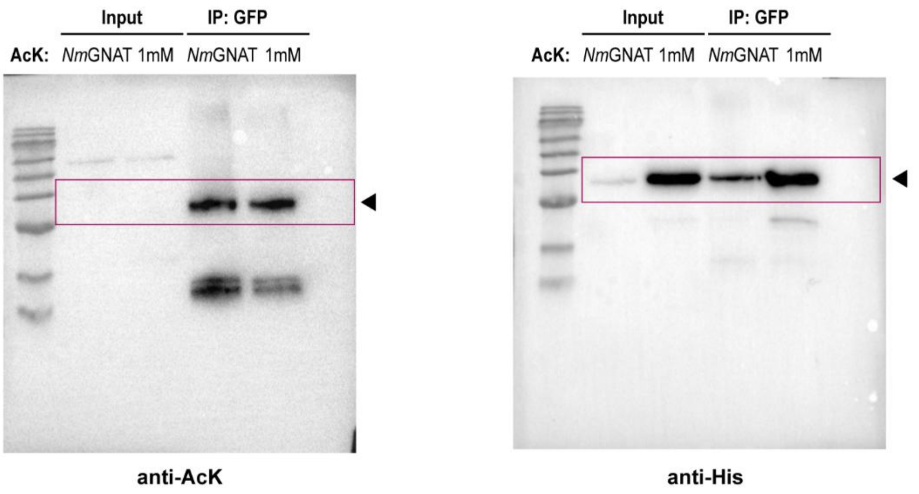
